# Single-particle analysis of small extracellular vesicles from the follicular fluid of women undergoing fertility treatments reveals distinct PD-L1^+^ populations

**DOI:** 10.1101/2024.12.17.628903

**Authors:** Barbara Bortot, Roberta Di Florio, Gabriella Zito, Francesco Valle, Marco Brucale, Giuseppe Ricci, Paola Viganò, Stefania Biffi

**Author notes:** Corresponding author Stefania Biffi, Via dell’Istria 65/1, 34137 Trieste, Italy.

## Abstract

In some cell systems, small extracellular vesicles bearing PD-L1 (PD-L1^+^ sEVs) have been shown to be able to suppress T-cell immunity. We have herein investigated whether a distinct profile of PD-L1+ sEVs exists in human follicular fluid (FF). Single-particle interferometric reflectance imaging sensing combined with a single-particle antibody capture and immunofluorescence labelling were used to determine the expression and colocalization of CD63, CD81, CD9, and PD-L1 in sEVs derived from FF of women undergoing fertility treatments (n=10). In addition, the size distribution of sEVs was investigated via atomic force microscopy. Our data indicate that the bulk of tetraspanin-expressing EVs in human FF are less than 50 nm in size. Tetraspanins and PD-L1 exhibit distinct expression and colocalization profiles at sEV level across all cohort samples. A total of 42%, 46%, and 50% of all the particles captured by anti-CD63, anti-CD81, and anti-CD9 antibodies, respectively, were positive for CD81. PD-L1 was expressed at the highest level on CD9+ sEVs, with an average value of 5% within the cohort. The presence of distinct PD-L1+ sEV subpopulations suggests that they may play a role in regulating the immune response in the follicular microenvironment. Further research is needed to fully understand the functional significance of PD-L1+ sEVs in this context and their potential as biomarkers for predicting fertility outcomes.

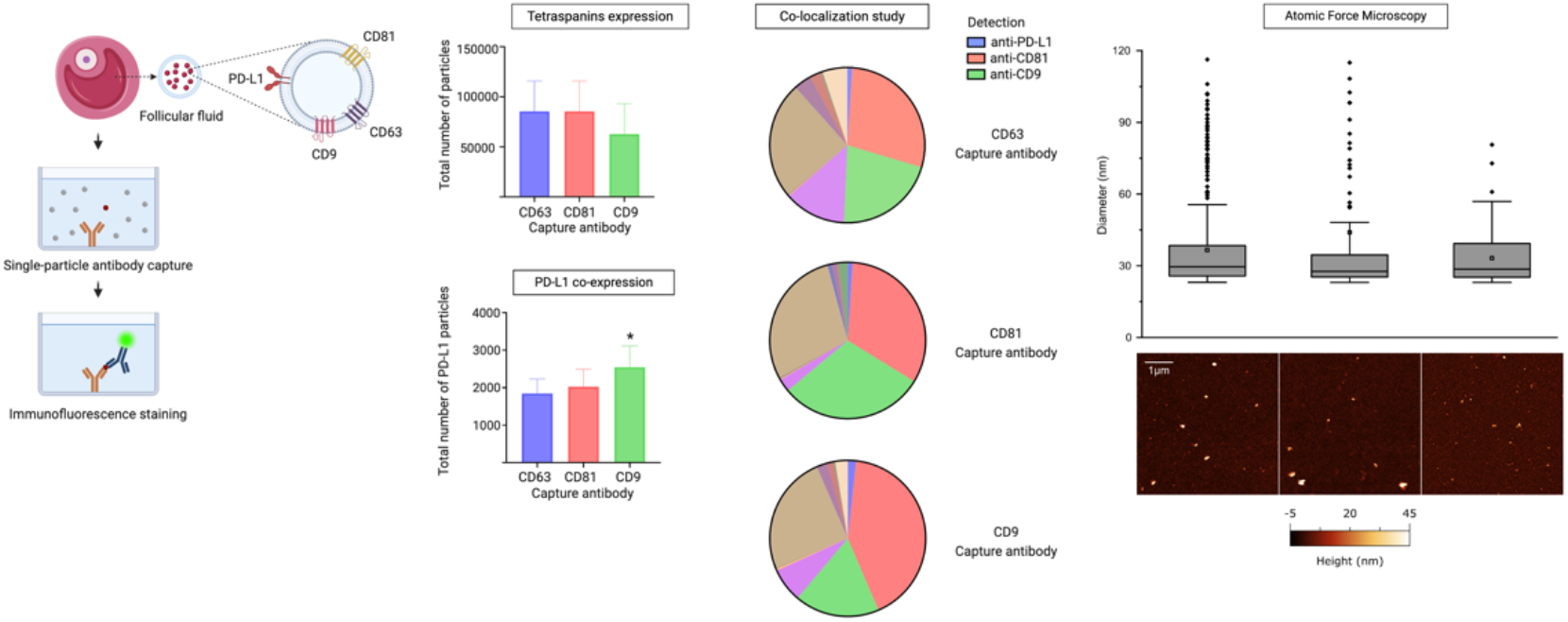

## Introduction

The follicular fluid (FF) surrounds developing oocytes in the ovaries, and its composition can significantly affect the success of fertilization and implantation (Dumesic et al., 2015; Revelli et al., 2009). FF is composed of active substances derived from the blood and secreted by granulosa and theca cells (Edwards, 1974; Fortune, 1994). It contains a variety of hormones, proteins, amino acids, enzymes, fatty acids, cytokines, blood-clotting factors, and other molecules that play essential roles in supporting follicle maturation (Da Broi et al., 2018; Saint-Dizier et al., 2019). Changes in FF composition reflect variations in the secretion of granulosa and theca cells, the inner membrane, and blood plasma, influenced by both normal and abnormal physiological processes (Edwards, 1974).

Extracellular vesicles (EVs) in FF have a distinctive composition and carry cargoes of messenger RNAs (mRNAs), miRNAs, proteins, and metabolites (Théry et al., 2018). These particles are actively released into the extracellular space and play an important role in cell-to-cell communication by delivering their cargo to target cells (Muraoka et al., 2024; Salomon et al., 2022; Tkach and Théry, 2016). Several studies have focused on the functional significance of EVs collected in FF in promoting cell-to-cell communication during the reproductive process (Salehi et al., 2023; Tesfaye et al., 2020; Wang et al., 2021). For example, Wang et al. (2021) studied noncoding RNAs present in small EVs (sEVs) in FF as potential pregnancy predictors, offering new avenues for utilizing EVs in assisted reproduction. In another study, the increase in the number of small EVs in the FF from older women has been correlated to an enhanced release from somatic follicular cells, likely controlled by TP53 signalling pathways (Battaglia et al., 2020). Additionally, six miRNAs were discovered to be upregulated in the follicular fluid of older women (Battaglia et al., 2020).

Recent research has shown that immune cells residing in the ovaries and fallopian tubes, such as T cells and macrophages, as well as nonimmune cells like granulosa cells and oocytes, express PD-1 and its ligands PD-L1 and PD-L2 (Johnson et al., 2023, 2019). PD-1 and its soluble ligands have been found in high concentrations in sEV fractions in human FF (Luu et al., 2020), at levels capable of controlling T-cell PD-1 activation when taken up, suggesting a regulatory role in immune responses (Johnson et al., 2023). Additionally, studies have shown that PD-L1 on EVs can promote tumor immune evasion by inducing T-cell exhaustion and inhibiting T-cell proliferation, highlighting PD-L1’s role in immune escape mechanisms employed by cancer cells (Chen et al., 2018). Therefore, activation of the PD-1 checkpoint pathway may be crucial for normal ovarian activities, such as follicle development and the establishment of immunological tolerance for germline cells and embryos (Johnson et al., 2023).

In order to investigate the intrafollicular communication mechanisms that regulate T lymphocyte immunity, the purpose of this study was to assess whether a distinct profile of PD-L1+ sEVs exists in human FF. This aim has relevance in the context of the recent demonstration for a role of PD-L1+ sEVs beyond tumor immunity (Yu et al., 2022).

## Materials and methods

### Patient recruitment

Ten patients who underwent assisted reproductive technology (ART) treatment were enrolled in this study (**Table 1**). Following approval by the Institutional Review Board (IRB-BURLO 07/2023 dd. 31.08.2023), patients were asked to sign an informed consent form. The FF from the first and largest punctured follicle from the bilateral ovaries was collected during the oocyte retrieval procedure (transvaginal follicular aspiration). Each ovarian follicle was aspirated independently.

**Table 1.** Patient characteristics.

| *Pt* | *Age* | *E2\**<br>*pg/ml* | *P4\**<br>*ng/ml* | *Final trigger* | *N*<br>*oocytes* | *Endometriosis* | *Pregnancy* | *Live birth* |
| --- | --- | --- | --- | --- | --- | --- | --- | --- |
| **1** | 33 | 324 | 2 | UP- hCG 10000 UI (Gonasi 10000 ®) | 24 |  | Yes** | yes |
| **2** | 34 | 2945 | 0,75 | UP- hCG 3300 UI (Gonasi 3300 ®) | 9 |  | Yes ** | yes |
| **3** | 35 | 1203 | 0,6 | UP- hCG 10000 UI (Gonasi 10000 ®) | 4 | yes | No |  |
| **4** | 41 | 683 | 0,7 | r-hCG 6500 UI (Ovitrelle250 ®) | 2 |  | No |  |
| **5** | 40 | 1371 | 1 | r- hCG 6500 UI (Ovitrelle250 ®) | 2 |  | No |  |
| **6 #** | 33 | 572 |  | 0.2 mg triptorelin (Decapeptyl ® ) | 20 |  | No |  |
| **7** | 43 | 2288 | 1,3 | r-hCG 6500 UI (Ovitrelle250 ®) | 3 |  | No |  |
| **8** | 31 | 2000 | 0,9 | 0.2 mg triptorelin (Decapeptyl ® ) | 6 |  | Yes** | yes |
| **9** | 35 | 2574 | 1,7 | HP-hCG 10000 UI (Gonasi 10000 ®) | 16 | yes | Yes** | No |
| **10** | 39 | 3172 | 08 | r-hCG 6500 UI (Ovitrelle250 ®) | 7 |  | No ** |  |

### Single-Particle Interferometric Reflectance Imaging Sensing: ExoView R100 Analysis

The investigation was performed via single-particle interferometric reflectance imaging sensing (SP-IRIS) via the ExoView R100 system and the ExoView Human Tetraspanin Kit (NanoView Biosciences, Brighton, MA, USA) (Mizenko et al., 2021). Samples were placed on an ExoView Tetraspanin Chip and incubated for 16 hours at room temperature. After that, they were washed three times with solution A from the ExoView Human Tetraspanin Kit from NanoView Biosciences. The immunocapture antibodies (anti-CD9 CF488, anti-CD81 CF555, and anti-CD63 CF647) were utilized in a diluted form at a concentration of 1:500 in solution A. To obtain the desired concentration of antibodies, 250 µL of the antibody solution was combined with the remaining 250 µL of solution A after washing the chip. This resulted in a final dilution of the antibodies at a ratio of 1:1000 for the incubation process. Following a 60-minute incubation at ambient temperature, the chips were cleansed and dehydrated, and images were captured via an ExoView R100 reader and ExoView Scanner 3.0 collection software. The acquired data were then examined via ExoView Analyser 3.0.

### Size exclusion chromatography isolation on qEVoriginal 35 nm columns, IZON

sEVs were isolated via size exclusion chromatography (SEC) according to the manufacturer’s instructions. Briefly, single qEV original 35 nm columns (Izon Science, Shaefer Italy) were allowed to reach room temperature and equilibrated with 17 ml of freshly filtered (22 μm) PBS. FF samples were first centrifuged at 2000 x g for 30 min. The supernatant was added at the top of the column (500 μl). After the sample was allowed to run into the column, 0.22-µm-filtered PBS was added to the top of the column tube. The following fractions were collected: F0 (1.6 mL = void volume of the column) and F1 to F6 (400 µL each).

Fractions F1-F6 were concentrated via ultracentrifugation (100,000 x g for 70 min).

### Atomic force microscopy (AFM) morphometry

Samples obtained via SEC isolation were analysed via AFM as described elsewhere (Bortot et al., 2022; Ridolfi et al., 2023, 2020). Briefly, images were taken in PeakForce mode on a Bruker Multimode8 equipped with a Nanoscope V controller, a sealed fluid cell and a type JV piezoelectric scanner using Bruker ScanAsystFluid+ probes (triangular cantilever, nominal tip curvature radius 2–12 nm, nominal elastic constant 0.7 N/m) calibrated with the thermal noise method. Image analysis was performed with a combination of Gwyddion 2.58 (Nečas and Klapetek, 2012) and custom Python scripts to recover the equivalent spherical diameter of individual particles.

## Results

### ExoView R100 analysis reveals the presence of sEVs with a distinct pattern of PD-L1 and tetraspanin profiles in FF

A comprehensive characterization of the tetraspanin expression profile in FF sEVs has not been accomplished; hence, we performed this investigation using a platform that minimized biases derived from the sEV isolation step and was capable of characterizing vesicles smaller than 50 nm (Mizenko et al., 2021). We have associated the analysis of tetraspanins with that of our target marker, PD-L1. The ExoView R100 platform was used to quantify distinct EV populations by combining two complementary systems: antigenic capture via interferometric imaging and fluorescent antigenic detection of the captured EV. Single-particle interferometric reflectance imaging (IM) enables the quantification of particles bound to the capture spot between 50 and 200 nm (Avci et al., 2015), whereas fluorescence mode was implemented to quantify the particles as small as 35 nm. The concurrent use of three distinct fluorescence-labelled antibodies enabled the assessment of antigen distribution on a single sEV (**Figure 1A**). The majority of sEVs were less than 50 nm in size, with anti-CD63 and anti-CD81 antibodies capturing the same number of EVs (37%) and anti-CD9 antibodies capturing fewer (27%) (**Figure 1B** and **1C**). Colocalization analysis revealed distinct patterns of EV populations (**Figure 1D**). EVs exhibited consistent patterns across the cohort of FF samples, with anti-CD81+ EVs representing the highest percentage at all capture locations. In particular, 42%, 46%, and 50% of all the particles captured by anti-CD63, anti-CD81, and anti-CD9 antibodies, respectively, were positive for CD81. The majority of the EVs colocalized with CD81 and CD9, indicating the presence of distinct populations of EVs that carry these proteins. A small proportion of EVs exhibited PD-L1 expression on the CD9, CD63, and CD81 antibody capture spots.

**Figure 1.**
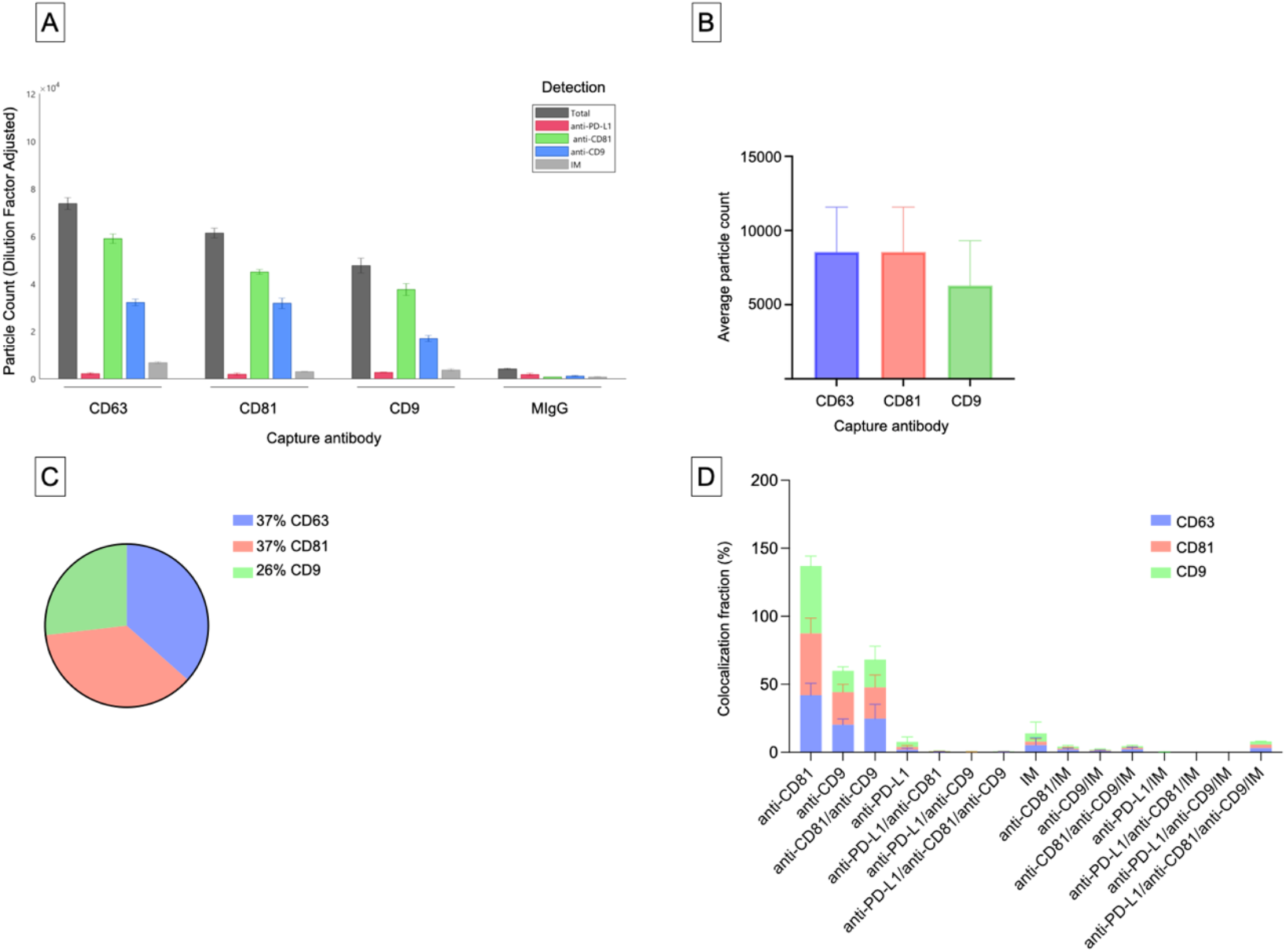
Tetraspanin marker expression profiles of individual sEVs. (**A**) Representative example of the tetraspanin colocalization pattern of follicular fluid (FF) obtained via specific antibodies (anti-CD63, anti-CD81, and anti-CD9) for capturing EVs. The captured EVs were then labelled with fluorescent anti-CD81, anti-CD9, and anti-PD-L1 antibodies. The population of EVs was visualized in ExoView fluorescence mode. Single-particle interferometric reflectance imaging (IM) was used to quantify the particles bound to the capture spot between 50 and 200 nm, whereas fluorescence mode was used to quantify the particles as small as 35 nm. (**B**) Average particle counts on the CD9, CD63, and CD81 antibody capture spots. (**C**) Particle fractions on the CD9, CD63, and CD81 antibody capture spots. (**D**) Protein colocalization analysis of the FF sample cohort was performed on the basis of single-EV protein expression data. Data are presented as percentages of the total fluorescence counts that correspond to each colour, indicating the expression combination of each protein.

### Characterization of PD-L1 expression profile of sEVs in FF

The analysis of the PD-L1 expression profile revealed a preference for colocalization with CD9+ sEVs (**Figure 2**). In terms of the three capture spots, CD63, CD81, and CD9, the percentages of EVs expressing PD-L1 were 2%, 3%, and 5%, respectively (**Figure 2A**). We observed that PD-L1 mostly colocalized with a single tetraspanin, with percentages of 2%, 2% and 4%, respectively, while the percentage of EVs exhibiting PD-L1 in combination with multiple tetraspanins was less than 1% (**Figure 2B**).

**Figure 2.**
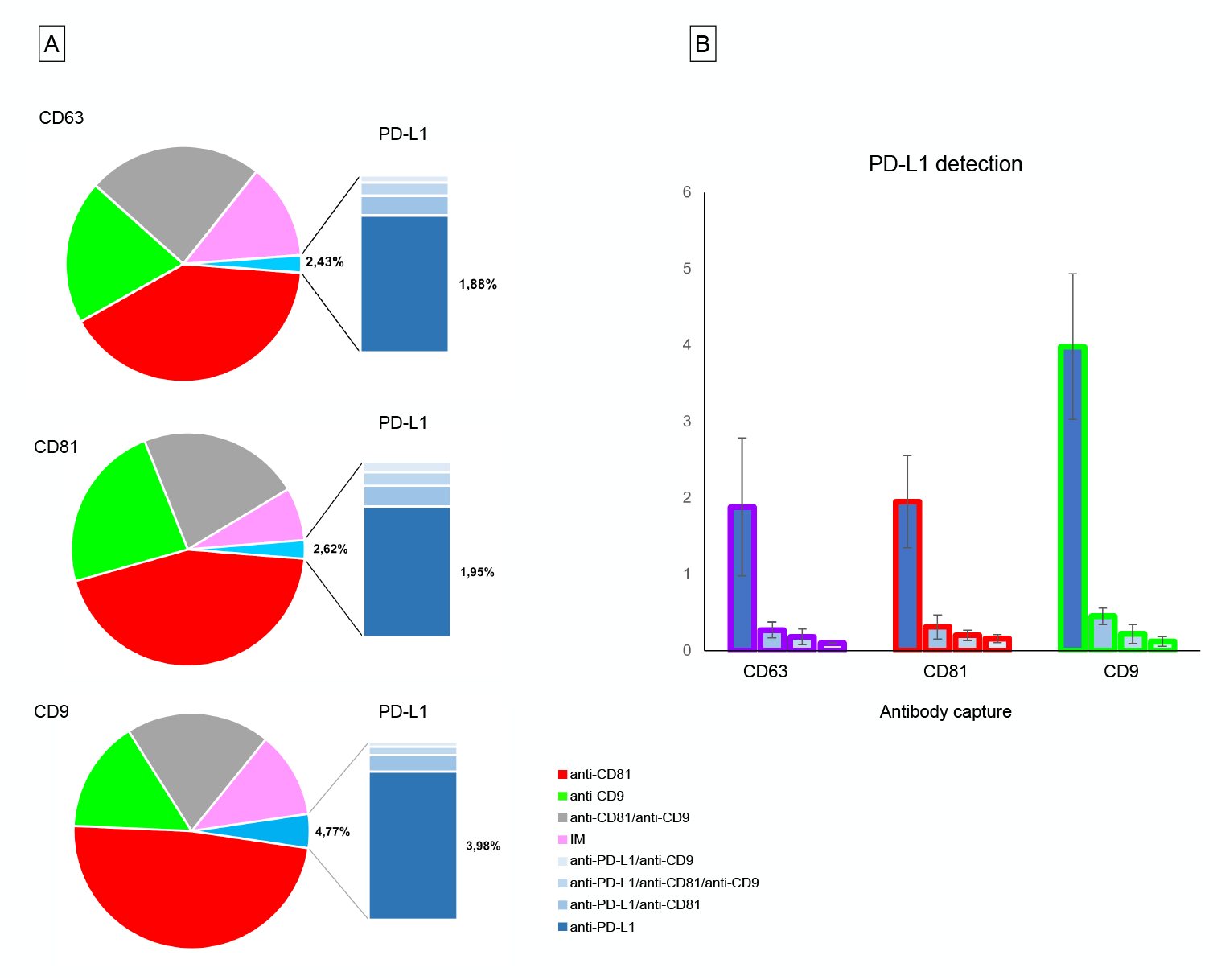
PD-L1 expression profiles of individual sEVs. (**A**) Protein colocalization analysis of the FF sample cohort on the CD9, CD63, and CD81 antibody capture spots. The bars in the pie charts show the percentages of PD-L1 + sEV subpopulations with different tetraspanin combinations. (**B**) Percentage of EVs expressing PD-L1 in combination with single or multiple tetraspanins on the CD63, CD81 and CD9 antibody capture spots.

It has recently been demonstrated that EVs exhibit a distinct nanomechanical fingerprint that can be identified using force spectroscopy. This fingerprint can be used to distinguish between various EV populations, making atomic force microscopy (AFM)-based nanomechanics a useful tool for evaluating EV identity, purity, and function (Bortot et al., 2021; Ridolfi et al., 2023). Single-particle AFM morphometry performed on SEC-enriched sEV samples revealed that over 50% of the particles had diameters smaller than 50 nm, while no significant amounts of vesicles with diameters above 100 nm were observed (**Figure 3**).

**Figure 3.**
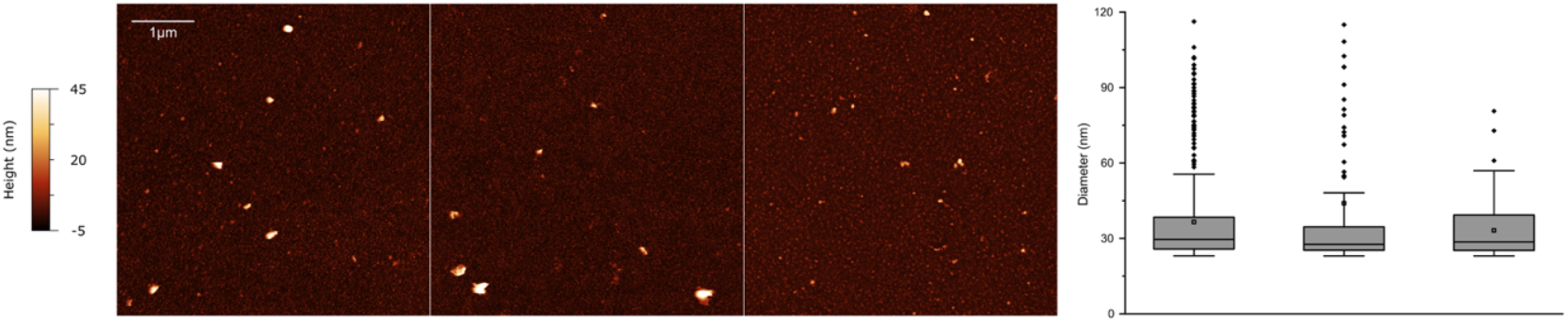
Representative AFM micrographs of three sEV samples from the cohort (left) and corresponding size distributions (right) obtained via quantitative morphometry on multiple images per sample. In all cases, the majority of particles have diameters below 50 nm.

## Discussion

EVs have emerged as important mediators and regulate several reproductive processes, including gamete development and implantation (Giacomini et al., 2017; Makieva et al., 2024; Smith and Russell, 2022). A growing body of evidence supports the idea that FF sEVs ensure the successful development of follicles and oocytes (Gonzalez Fernandez et al., 2023; Tesfaye et al., 2020; Wang et al., 2021). Furthermore, adverse pregnancy outcomes in ART have been linked to the low quality of embryos (Fauque et al., 2007), with FF sEVs being potential non-invasive biomarkers for predicting pregnancy (Muraoka et al., 2024). However, EV characterization in human FF is still in its early stages, and more research in this area is needed to fully understand the potential of EVs in FF and their impact on female reproductive biology (Giacomini et al., 2020).

On the basis of our data, most tetraspanin-expressing EVs in human FF are smaller than 50 nm. This contrasts with previous studies on human samples. Several studies have successfully isolated FF sEVs via serial centrifugation or precipitation and revealed that sEVs are less than 200 nm in diameter, with a mean size of approximately 100 nm (Gu et al., 2024; Muraoka et al., 2024; Zhou et al., 2024). The different techniques used for EV analysis may account for this difference. The strengths of the present findings are that we followed an innovative approach to identify and characterize EVs from unpurified biological sources. The Exoview R100 platform allows direct capture of EVs from biofluids via tetraspanin-specific antibodies, eliminating the need for prior separation. Moreover, SP-IRIS on the ExoView R100 platform enables the detection of EVs smaller than 50 nm, a capability that other methods such as nanoparticle tracking analysis (NTA) cannot achieve (Bachurski et al., 2019). AFM analysis of sEVs at the single-vesicle level revealed their size distribution, confirming that most sEVs are under 50 nm in size (Ridolfi et al., 2023, 2020).

The tetraspanins CD9, CD63, and CD81 are widely recognized as the most prevalent markers associated with EVs in the scientific literature (Karimi et al., 2022; Okada-Tsuchioka et al., 2022). These markers have been extensively used in various research investigations, such as ELISA, flow cytometry, and lab-on-a-chip assays, to capture EVs comprehensively. We observed consistent patterns in the EV subpopulations based on tetraspanins, with anti-CD81+ sEVs representing the majority of sEVs in all samples and antibody-capture spots. In particular, 42%, 46%, and 50% of all the particles captured by anti-CD63, anti-CD81, and anti-CD9 antibodies, respectively, were positive for CD81. These results are consistent with prior studies linking CD81 and CD9 in the context of reproductive biology (Kaji et al., 2002; Takahashi et al., 2001). In particular, Rubinstein and colleagues studied the fertility of mice lacking the CD9 and CD81 genes suggesting that CD9 and CD81 are crucial for successful sperm-egg interactions (Rubinstein et al., 2006).

CD9+ sEVs have the highest PD-L1 expression at 5%. Additionally, we observed that PD-L1 was predominantly associated with a single tetraspanin protein rather than multiple tetraspanins in sEVs. The tetraspanin CD9 is expressed by all the major leukocyte subsets, including B cells, CD4+ T cells, CD8+ T cells, natural killer cells, granulocytes, monocytes, macrophages, and both immature and mature dendritic cells, as well as at elevated levels by endothelial cells (Reyes et al., 2018). Notably, CD9 mediates the interaction between sperm and eggs during fertilization (Chalbi et al., 2014; Kaji et al., 2000; Le Naour et al., 2000; Miyado et al., 2000; Rubinstein et al., 2006), allowing their membranes to fuse for successful fertilization. The fact that PD-L1 expression is much greater in the CD9+ sEV population, which is actually less abundant than the CD63+ and CD81+ subsets, suggests that CD9 might play a particular role in delivering PD-L1. A weakness of the present study relates to the limited number of FF samples analysed. On the other hand, results were consistent among samples as demonstrated by the narrow range of variability in the data.

In conclusion, the presence of PD-L1 in FF sEVs suggests that they may be involved in maintaining immune homeostasis within the ovarian microenvironment through the modulation of EV-mediated signalling pathways. Although sample size does not allow for correlation analysis between protein expression on sEVs and clinical parameters or pregnancy outcomes, these findings highlight the importance of further research into the role of the PD-1 checkpoint pathway in female reproduction.

